# *Aspergillus* species in Iceland: species distribution and *cyp51A* mutation testing in a 7.5-year clinical isolate collection

**DOI:** 10.1101/2024.08.08.607273

**Authors:** Ingibjorg Hilmarsdottir, Ervin Martien Alcanzo, Bert Gerrits van den Ende, Jos Houbraken, Ferry Hagen

## Abstract

**Introduction.:** Among challenges in diagnosing and treating aspergillosis are *Aspergillus* species identification, which can inform therapeutic choices, and detection of antifungal resistance. Increasing rates of triazole resistance have been reported for the most common species, *Aspergillus fumigatus*.

**Methods.:** A collection of clinical *Aspergillus* isolates cultured during 2014–2021, previously identified morphologically, were tested with the AsperGenius qPCR-kit for identification of *A. fumigatus* and *Aspergillus flavus* and the presence of a *cyp51A* mutation encoding azole resistance (TR_34_/L98H and Y121F/T289A); other species were identified with calmodulin sequencing.

**Results.:** Identification of the 312 isolates (224 patients) tested revealed 19 species in nine *Aspergillus* sections. Rare or cryptic species were identified in 48 isolates (44 patients). Thirty-one isolates had been assigned an incorrect species name when initially cultured. The TR_34_/L98H and Y121F/T289A mutations were not found in any of the 202 *A. fumigatus* isolates tested.

**Discussion.:** In this first study on clinical *Aspergillus* isolates in Iceland the proportion of cryptic and rare species was on par with findings from other countries. In contrast, unlike other studies we did not find the most common azole resistance-related mutations in *A. fumigatus*, possibly due to the absence of significant environmental antifungal pressure in the local agriculture.

## Introduction

*Aspergillus* species are ubiquitous environmental molds responsible for chronic and acute, life-threatening infections in humans, mainly in the respiratory tract. First line treatment options include amphotericin B, whose use is limited by toxic side-effects, and triazole agents which are generally well tolerated.^1^ However, increasing rates of resistance to triazoles have been reported for the most common species, *Aspergillus fumigatus*, which is now categorized as a priority fungal pathogen for research and public health efforts by the World Health Organization.^2^

Some of the challenges in diagnosing and treating aspergillosis involve *Aspergillus* species identification—which can inform therapeutic choices—and detection of antifungal resistance.

Among the more than 100 *Aspergillus* species reported in clinical specimens many are rare or cryptic species for which conventional identification methods, based on morphological characteristics, may be unreliable.^3,4^ While *A. fumigatus* remains the most common species in deep-seated aspergillosis, molecular methods have revealed cryptic *Aspergillus* species in 3–30% of isolates from patient specimens in the USA, Europe and East Asia.^4–8^

Both common and rare or cryptic species may show decreased antifungal susceptibility which highlights the importance of timely species identification and access to susceptibility testing.^6,9–11^ Though *A. fumigatus* is generally susceptible to triazole agents, recent studies from all inhabited continents have revealed resistance rates in the range of 1–14% for clinical and environmental isolates, with a few European centers reporting up to 30% for some patient groups.^10^ Triazole resistance in *A. fumigatus* is usually due interruption of ergosterol synthesis in the cell membrane secondary to changes affecting the *cyp51A* gene which encodes 14α-lanosterol demethylase.^10^ Most commonly the changes involve tandem repeats in the *cyp51A* promoter and point mutations in the *cyp51A* gene, i.e. TR_34_/L98H and TR_46_/Y121F/T289A, which are believed to be driven in large part by agricultural use of triazole agents for various crops.^10,12^ Treatment choices for aspergillosis are further complicated by the fact that some cryptic species, e.g. *Aspergillus lentulus* and *Aspergillus calidoustus*, show high minimal inhibitory concentration values for triazoles.^9,11,13,14^

The aim of this study was to provide the first report on *Aspergillus* species distribution and triazole resistance in *A. fumigatus* from clinical specimens in Iceland, a geographically isolated country with negligible crop farming and low agricultural use of fungicides.

## Methods

### Setting and Aspergillus isolates

The Department of Microbiology at Landspítali University Hospital, a 650-bed institution, serves as the primary diagnostic laboratory for more than 60% of the roughly 390,000 population in Iceland and is the reference center for the identification of all filamentous fungi isolated from clinical specimens nationwide. *Aspergillus* species are identified using conventional methods based on macroscopic and microscopic morphology features. Cryopreservation of *Aspergillus* species (on cryobeads at -80°C) isolated from paranasal sinuses, the lower respiratory tract and normally sterile tissues was implemented in mid-2014. Laboratory procedures prescribe preserving *Aspergillus* species isolates cultured for the first time from a patient and repeat isolates of *A. fumigatus* if cultured more than a week from the previous isolate. In addition, starting in early 2019 *Aspergillus* species isolates from other specimen types have been randomly selected for preservation on cryobeads at - 80°C.

The laboratory information system was searched for *Aspergillus* species for the period of July 2014 to December 2021. A total 722 *Aspergillus* isolates were cultured from 699 specimens belonging to 479 patients. The specimens included 408 from the respiratory tract, 163 from keratinized tissue, mostly nail scrapings, 102 from ears and 26 from various sites. Morphological identification yielded *A. fumigatus* in 374 isolates (226 patients), *Aspergillus niger* in 116 (102 patients), *Aspergillus flavus* in 107 (66 patients), unidentified *Aspergillus* species in 91 (89 patients and mostly keratinized specimens), *Aspergillus terreus* in 25 (22 patients) and various species in nine. Cryobeads containing isolates that had been preserved during the period were sent to the Westerdijk Fungal Biodiversity Institute (Utrecht, The Netherlands) where *Aspergillus* species were revived on malt extract agar at 35°C and conidia were stored in tryptone soy broth at -80°C until further use.

### Species identification and detection of azole resistance

After subculturing, genomic DNA was extracted by using the DNAeasy® Ultraclean Microbial Kit (Qiagen, Hilden, Germany) according to manufacturer’s instructions. To separate *A. fumigatus* from other found species all samples were screened with the AsperGenius qPCR-kit (PathoNostics, Maastricht, Netherlands) targeting the multicopy 28S rRNA gene to detect *A. fumigatus*, *A. flavus* and pan-*Aspergillus*. For all *A. fumigatus* positive samples, the azole resistance assay of the AsperGenius qPCR-kit was applied which targets the TR_34_, L98H, Y121F, and T289A point mutations in the *cyp51A* locus.^15^

*Aspergillus* species that were not identified by the AsperGenius qPCR as *A. fumigatus* were subsequently characterized using partial sequencing of the calmodulin gene (*CaM*) as previously described.^16^

Sequences were generated on the ABI3730xL Genetic Analyzer platform (Applied Biosystems, Palo Alto, CA, USA), and raw data was manually checked and corrected using the program SeqMan Pro (Lasergene v17 software package; DNASTAR, Madison, WI, USA). The species identification process was performed using BLASTn on corrected sequences using the in-house curated *Aspergillus* calmodulin sequence database.^16^ Calmodulin sequences were deposited into the NCBI Genbank with accession numbers OQ181235– OQ181346.

### Ethics statement

The authors confirm that the ethical policies of the journal, as noted on the journal’s author guidelines page, have been adhered to and the appropriate ethical review committee approval has been received. The study was approved by the National Bioethics Committee of Iceland (license nr. 22-100).

## Results

Of the approximately 350 preserved *Aspergillus* isolates, 312, from 302 specimens belonging to 224 patients, were retrieved from freezers, screened with the AsperGenius qPCR-Kit and, when negative, subsequently identified by sequencing a part of the calmodulin gene (Table 1). Thirty patients had multiple specimens, two to 18 per patient, yielding a total of 114 isolates (Table 2). Among the 25 patients having multiple lower respiratory tract specimens, 15 had underlying chronic lung disease or cystic fibrosis and seven had underlying malignancy or other immunocompromizing conditions.

**Table 1.**
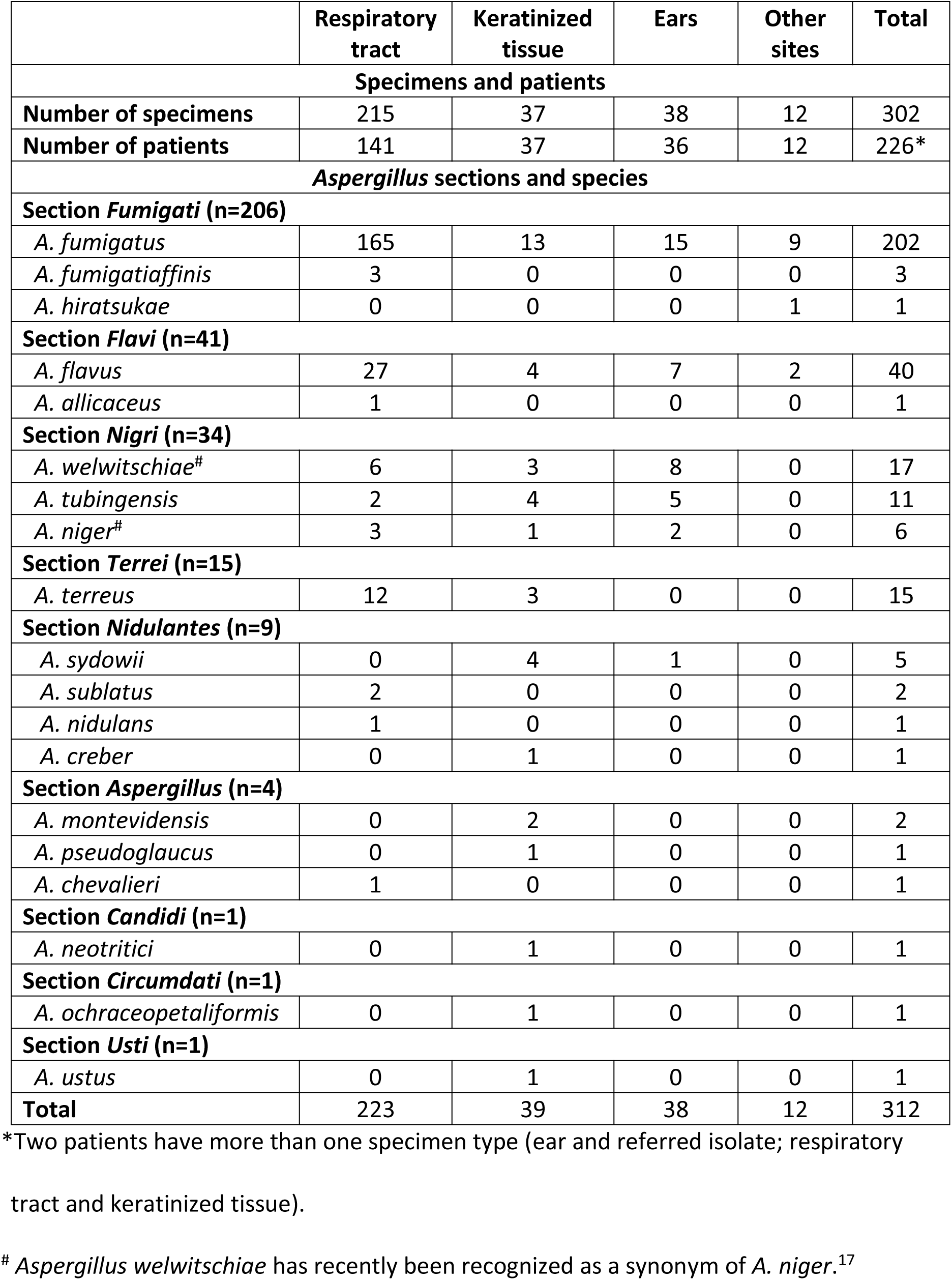
Specimen types and *Aspergillus* species (n=224 patients)

**Table 2.**
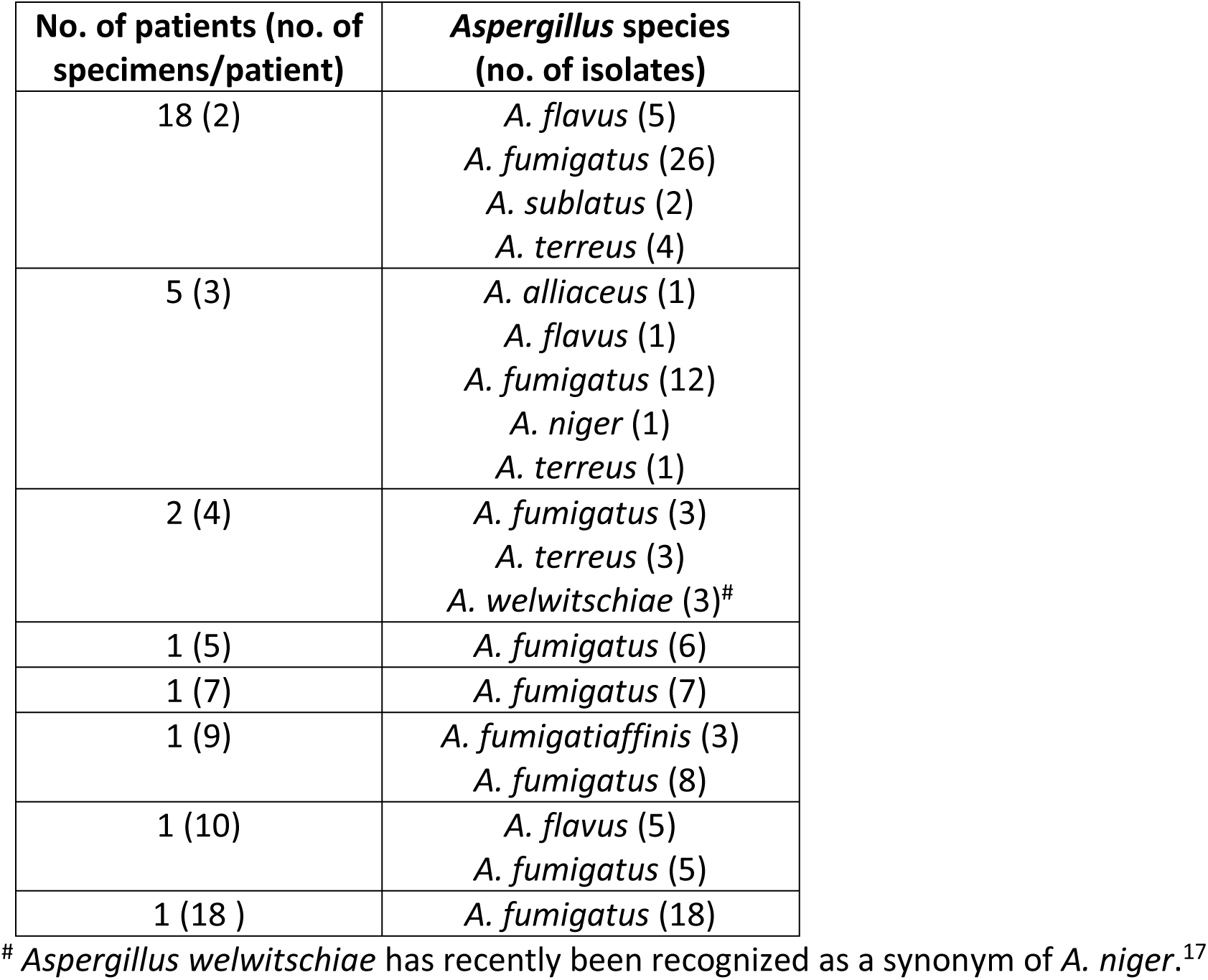
*Aspergillus* species from patients with multiple specimens (n=30 patients)

Among the 224 patients, 210 had one *Aspergillus* species and 14 had multiple species in the same or separate specimens. Results from AsperGenius qPCR and calmodulin sequencing revealed 19 *Aspergillus* species in nine sections (Table 1). Isolates belonged to section *Fumigati* in 144 patients, section *Flavi* in 35, section *Nigri* in 32, section *Terrei* in 12, section *Nidulantes* in nine and other sections in seven. Rare or cryptic *Aspergillus* species, i.e. other than *A. flavus*, *A. fumigatus*, *A. niger*, *A. nidulans* and *A. terreus*, were found in specimens from 44 patients and were more prevalent in superficial specimens (from 33 out of 78 patients) than in respiratory specimens (from 11 out of 141 patients,)

Misidentification occurred for 31 (from 28 patients) of the total 312 *Aspergillus* isolates studied. They involved cryptic and rare species that had been assigned to the correct *Aspergillus* section but incorrect species when morphologically identified. In addition 28 isolates had been identified to genus level only (Table 3).

**Table 3.** *Aspergillus* identification discrepancies.

| Morphological identification | Molecular identification | Number of isolates |
| --- | --- | --- |
| **Misidentification (n=31)** |  |  |
| *A. flavus* (section *Flavi*) | *A. alliaceus* * | 1 |
| *A. fumigatus* (section *Fumigati*) | *A. fumigatiaffinis* * | 2 |
| *A. nidulans* (section *Nidulantes*) | *A. sublatus* * | 1 |
| *A. niger* (section *Nigri*) | *A. tubingensis* * | 10 |
|  | *A. welwitschiae* ** | 17 |
| **Incomplete identification (n=28)** |  |  |
| ***Aspergillus* species** | *A. chevalieri* | 1 |
|  | *A. flavus* | 4 |
|  | *A. fumigatiaffinis* | 1 |
|  | *A. fumigatus* | 4 |
|  | *A. hiratsukae* | 1 |
|  | *A. montevidensis* | 2 |
|  | *A. nidulans* | 1 |
|  | *A. niger* | 1 |
|  | *A. ochraceopetaliformis* | 1 |
|  | *A. pseudoglaucus* | 1 |
|  | *A. creber* | 1 |
|  | *A. sublatus* | 1 |
|  | *A. sydowii* | 5 |
|  | *A. terreus* | 1 |
|  | *A. neotritici* | 1 |
|  | *A. tubingensis* | 1 |
|  | *A. ustus* | 1 |

\*Belong to the same *Aspergillus* section as the morphologically identified isolate(s).

Azole resistance-associated mutations in the *cyp51A* gene were not detected in any of the 202 *A. fumigatus* isolated tested with the AsperGenius qPCR assay.

## Discussion

In this first molecular study on clinical *Aspergillus* isolates in Iceland, most of which were from the respiratory tract, rare or cryptic species were identified in 15.4% of isolates (48/312) and 19.6% of patients (44/224). Common azole resistance-associated alterations in the *cyp51A* gene were not detected in any *A. fumigatus* isolate.

Whereas 98% of *A. fumigatus* (198/202), 93% of *A. terreus* (14/15) and 90% of *A. flavus* (36/40), were correctly identified to species level via morphological characteristics, all isolates of cryptic and rare species were assigned incorrect species name or identified to genus level only. Other investigators have illustrated the limitations of morphological identification^4^ and the importance of employing molecular methods^4–8^ for correct assignment of rare and cryptic species.

None of the 202 *A. fumigatus* isolates tested in our study was found positive for the most commonly detected azole resistance markers, TR_34_/L98H and Y121F/T289A (note that AsperGenius does not detect the TR_46_ associated with Y121F/T289A), which contrasts with findings from the majority of other Western European countries where these *cyp51A* gene alterations have been demonstrated in clinical, and sometimes also environmental, isolates.^18–22^

Evidence suggests TR_34_/L98H and TR_46_/Y121F/T289A emerged subsequent to the introduction of azole fungicides for crop agriculture in the 1970s.^10^ These agents constitute the largest group of sterol biosynthesis inhibitors which represent over 30% of globally used fungicides in agriculture.^23^ According to a recent Eurostat report, over 40% of pesticides, i.e. 150,000 tonnes, sold in Europe in 2021 were fungicides and bactericides and 7% of these were azole compounds.^24^ In Iceland, 650 kg of fungicides were imported in 2021 (none is produced locally)^24^ and during the seven preceding years the imports never exceeded half a ton (personal communication from The Environment Agency of Iceland). Only two azoles, penconazole and propiconazole, have been available in the country for the past decade, with imported amounts peaking at 60 kg in 2015 followed by a manifold decrease to zero in 2021 (personal communication from The Environment Agency of Iceland). Penconazole has been used mainly indoors, for greenhouse cultures, and propiconazole, whose sales have not been authorized in Europe^25^ since 2020, was used outdoors for cereal grain production and in greenhouses. Overall, Iceland appears to have the lowest pesticide sales among countries in the European Economic Area^24^, most likely explained by the country’s harsh climate and cool growing season, which limit the production of crops other than hay.^26,27^ In addition, the geographical isolation may prevent or delay transmission of plant pathogens to the island.

The above indicates that *A. fumigatus* populations present in Iceland are unlikely to acquire azole resistance mechanisms that are related to antifungal pressure in the environment. Azole-resistant strains could, however, be carried to the island via humans, imported goods, and possibly wind dispersal.^28,29^ Surveillance of azole resistance in our local *A. fumigatus* collection, as presented in this study, should therefore be conducted regularly, and such surveillance is indeed recommended by the ESCMID-ECMM-ERS guideline on the diagnosis and management of *Aspergillus* diseases.^30^

The main limitation to our study was that conventional antifungal susceptibility testing was not performed and we can therefore not rule out the presence of azole resistance in our isolates via mutations other than TR_34_/L98H and TR_46_/Y121F/T289A.

## Acknowledgements

We thank Helga Ösp Jónsdóttir at The Environment Agency in Iceland for information about fungicide sales. Pathonostics (Maastricht, The Netherlands) is acknowledged for providing the AsperGenius kits free-of-charge.

## Author Contributions

**Ingibjorg Hilmarsdottir:** conceptualization, funding acquisition, investigation, methodology, project administration, resources, supervision, validation, writing – original draft preparation, writing – review & editing. **Ervin M. Alcanzo:** data curation, formal analysis, investigation, validation, writing – review & editing. **Bert Gerrits van den Ende:** data curation, formal analysis, investigation, validation, writing – review & editing. **Jos Houbraken:** data curation, formal analysis, investigation, resources, validation, writing – review & editing. **Ferry Hagen:** conceptualization, funding acquisition, investigation, methodology, project administration, resources, supervision, validation, writing – original draft preparation, writing – review & editing.

